# MatchMaker: A Deep Learning Framework for Drug Synergy Prediction

**DOI:** 10.1101/2020.05.24.113241

**Authors:** Halil Ibrahim Kuru, Oznur Tastan, A. Ercument Cicek

**Affiliations:** Department of Computer Engineering, Bilkent University, Ankara, Turkey 06800; Faculty of Engineering and Natural Sciences, Sabanci University, Istanbul, Turkey 34956; Computational Biology Department, Carnegie Mellon University, Pittsburgh, PA 15213

**Keywords:** drug synergy prediction, neural networks, supervised learning, drug combinations

## Abstract

Drug combination therapies have been a viable strategy for the treatment of complex diseases such as cancer due to increased efficacy and reduced side effects. However, experimentally validating all possible combinations for synergistic interaction even with high-throughout screens is intractable due to vast combinatorial search space. Computational techniques can reduce the number of combinations to be evaluated experimentally by prioritizing promising candidates. We present MatchMaker that predicts drug synergy scores using drug chemical structure information and gene expression profiles of cell lines in a deep learning framework. For the first time, our model utilizes the largest known drug combination dataset to date, DrugComb. We compare the performance of MatchMaker with the state-of-the-art models and observe up to ~ 20% correlation and ~ 40% mean squared error (MSE) improvements over the next best method. We investigate the cell types and drug pairs that are relatively harder to predict and present novel candidate pairs. MatchMaker is built and available at https://github.com/tastanlab/matchmaker

## 1 Introduction

In complex diseases, multiple cellular mechanisms are often altered in the cell; therefore, treating them with a single drug and focusing on a single target is usually an ineffective strategy. Combination therapy is a promising solution that uses multiple drugs that exhibit better therapeutic efficacy together than the sum of individual drug effects [1]. One other advantage of combination therapy is that by lowering the required dosage per drug, it reduces the risk of adverse effects [2]. Especially, in cancer, where drug resistance is common, combinatorial drug therapy has been an effective strategy for several decades [3, 2, 4]. Thus, pinpointing the right combinations of drugs is an important task that has translational, clinical, and economic impacts.

It is possible to test for synergistic activity of drugs experimentally. However, exhaustive testing of drug combinations is not feasible even with high-throughput screens due to the vast number of drug combinations [5]. Computational methods that predict drug pair synergy can complement and guide high-throughput synergy screens by prioritizing the drug pairs to be further evaluated by experiments [2]. To this end, several algorithms have been proposed to predict synergistic drug pairs which utilize a diverse set of features such as chemical structure, biological networks interactions (e.g., drug-protein, protein – disease, etc.) and omics data [6, 7, 8, 9, 10, 11, 12, 13, 14, 15]. While feature engineering and a systems approach come with the promise of performance increase, features other than the chemical structure are not always available, and biological networks such as protein interaction networks are incomplete. This limits the generalization and applicability of such approaches.

Two algorithms achieve state-of-the-art performances on drug synergy prediction solely by using the drug structures and gene expression profile of the target cell. Preuer et al. propose a deep neural network based approach, DeepSynergy, which uses only the chemical fingerprints of the drugs and the gene expression profiles of the cell line of interest. Janizek et al. present TreeCombo, an algorithm based on the extreme gradient boosted trees (XGBoost) algorithm [18] on the same dataset with the same set of features and report performance improvements over DeepSynergy.

In this study, we present *MatchMaker*, a deep neural network-based drug synergy prediction algorithm. The model makes use of the drug chemical structures and untreated cell line gene expression profiles to predict the Loewe Additivity score [19] of a drug pair (referred to as Loewe score from now on). Unlike previous work [17, 16], which concatenate the features and predict synergy, MatchMaker trains two parallel subnetworks, one per drug, to learn drug specific representation on a particular cell line. The joint representation for the pair then becomes an input for a third sub-network, which learns to predict drug pair synergy. We show that this approach improves over state-of-the-art performance. Moreover, for the first time in the literature, we utilize the largest drug combination dataset released to this date [20] and benchmark the performances of the state-of-the-art methods on this dataset.

MatchMaker achieves 79.49 Mean Squared Error (MSE), 0.79 Pearson and 0.74 Spearman correlation when predicting Loewe scores. This corresponds to 42%, 16%, and 19% improvements over the state of the art, respectively. We investigate the cell lines and drug types for which the model is effective in predicting synergies. We detect novel synergistic pairs that are not marked in the ground truth data set but have reports in the literature as effective pairs. MatchMaker is built and available at https://github.com/tastanlab/matchmaker.

## 2 Methodology

### 2.1 Problem Formulation

We model the synergy prediction problem as a regression task. For a given drug pair < *i*, *j* > and a cell line *k*, target value is *y*_*i,j,k*_ ∈ ℝ which is measured by Loewe [19] and positive *y*_*i,j,k*_ values indicate synergism. Our dataset contains *n* unique trios. Each trio contains two drugs *i* and *j* represented with *x*_*i*_ and *x*_*j*_ drug feature vectors. The cell line *k* is represented with *c*_*k*_ cell line feature vector. The problem is to predict *y*_*i,j,k*_ by using drug (*x*_*i*_ and *x*_*j*_) and cell line specific features (*c*_*k*_).

### 2.2 Drug Combination Dataset

We use the synergy scores provided in the DrugComb [20] database. The data were obtained from https://drugcomb.fimm.fi/ (version v1.4, downloaded on Dec 2019). DrugComb curates high-throughput combinatorial drug screening data from four studies: (i) O’Neil dataset [21], (ii) Forcina dataset [22], (iii) NCI Almanac dataset [23], and (iv) Cloud dataset [24].

The dataset contains 466, 033 combinations across 112 cell lines and 4, 150 drugs. Since we use drug structure and cell line gene expression together as features, we filter this data such that the drugs with the available structure information from PubChem database [25] and cell lines with available gene expression data from Iorio et al.. This leaves us with 335, 692 drug pair-cell line combinations that cover 3, 040 drugs and 81 cell lines. For the 81% of the combinations, there is only a single experiment reporting Loewe score. For the remaining combinations, there are more than one (2 to 8) replicate experiments reporting Loewe scores. For some of these cases, the replicate experiments disagree. That is, some replicates have an equal number of positive Loewe scores (synergism) and negative scores (antagonism) on drug pair, cell line trio. We filter out such 1, 861 trios, in which half of the replicates report synergism and other half reports antagonism. For other trios, the Loewe scores of the replicate experiments are averaged. The final processed data contains 286, 421 experiments (drug1, drug2, cell line) for 3, 040 drugs, performed on 81 cell lines. We should note that not all drug pairs are tested on every cell line, and on average, a drug pair is tested on 25 cell lines. The Loewe distribution of the scores is shown in Figure 1.

**Fig. 1:**
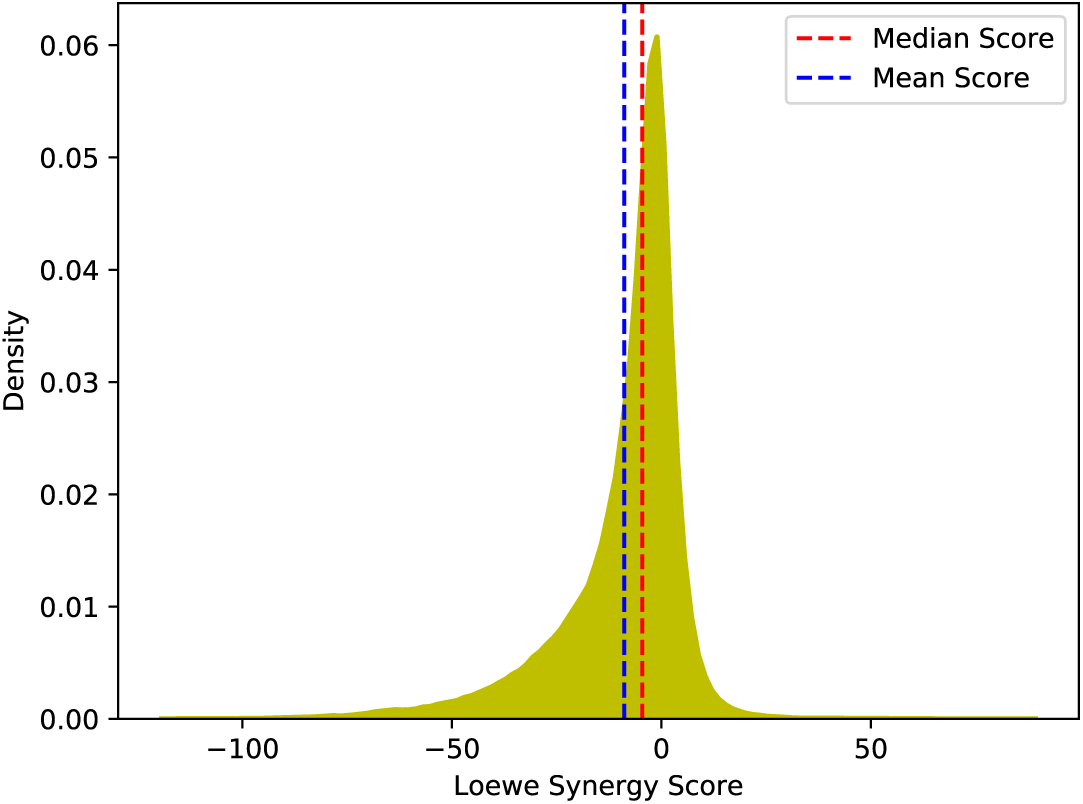
The distribution of the Loewe scores in the processed DrugComb dataset. The processed data contains 286, 421 experiments (drug1, drug2, cell line) for 3, 040 drugs, performed on 81 cell lines.

### 2.3 Input Features

We use two types of features to train the model. The first set of features encodes the untreated cells’ gene expression (GEX) profiles. The data were obtained from Iorio et al. (accession number E-MTAB-3610 at ArrayExpress [27]). We downloaded the RMA normalized gene expression profiles from https://www.cancerrxgene.org (downloaded on Dec 2019). For each cell line *k*, we use the expression profiles of landmark genes, denoted as *c*_*k*_ ∈ ℝ^972^. We obtain the list of landmark genes from Subramanian et al.. The second set of features describes the drug pairs’ chemical structure. To represent the drugs in the combination, drug chemical descriptor features *x* ∈ ℝ^541^ are calculated using *ChemoPy* Python library [29]. The chemical descriptors and the number of features for each descriptor are provided in Table S1.

### 2.4 The MatchMaker Architecture

MatchMaker is a deep neural network-based drug synergy prediction algorithm. The architecture is shown Figure 2. The model inputs (i) the drug chemical descriptors of a pair of drugs and (ii) an untreated cell line gene expression profile as features to predict the Loewe score [19] of a drug pair. The model contains three neural subnetworks. First, there are two subnetworks that learn a representation of each drug conditioned on cell line gene expression of the given cell line. The output of these two subnetworks are then concatenated and input to a third subnetwork that predicts the final output. The individual components are detailed below.

**Fig. 2:**
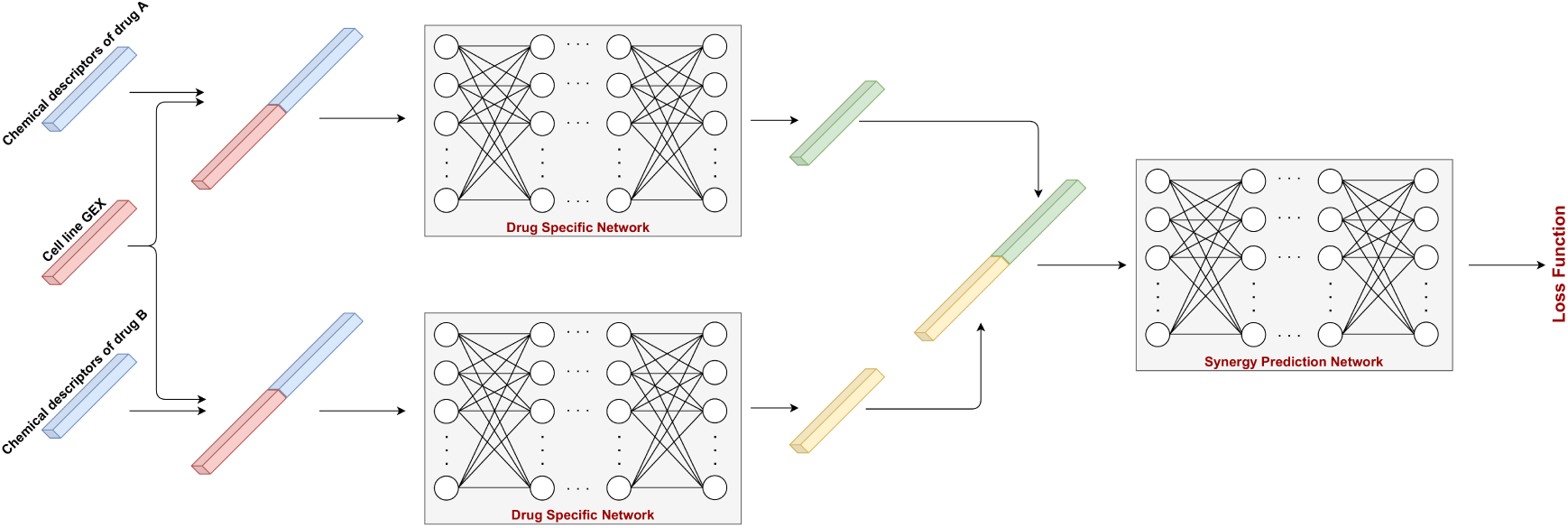
The architecture of MatchMaker. The chemical features of each drug are concatenated with gene expression features of the cell line and input to drug specific subnetworks (DSNs). The DSNs learn a representation of each drug separately conditioned on cell line gene expressions. Synergy Prediction Network (SPN) takes the representations of both drugs and applies fully connected layers to predict Loewe values. The model is trained to minimize a weighted version of the MSE between actual and predicted values in an end-to-end fashion.

#### Drug Specific Subnetworks (DSN)

For a drug pair, cell line trio, MatchMaker includes two Drug Specific Subnetworks, *DSN*_*i*_, and *DSN*_*j*_ for each drug *i* and *j* respectively. *DSN*_*i*_ inputs the i^*th*^ drug’s chemical representation (*x*_*i*_) together with untreated gene expression profile of the *k*^*th*^ cell line (*c*_*k*_). Symmetrically, *DSN*_*j*_ inputs *x*_*j*_ and *c*_*k*_. Both DSNs share the same architecture and apply three fully connected (FC) layers, followed by activation functions and dropout [30]. We use ReLU activation [31] at the first two FC layers and linear activation at the final layer of DSNs. Dropout probabilities are set to 0.2 and 0.5 at the first and second layers, respectively. No dropout is applied for the final layer. Each DSN produces a vector embedding for the corresponding drug conditioned on the cell line expression. We refer these embeddings as 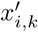 and 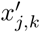. The formulation of DSN can be summarized as follows:

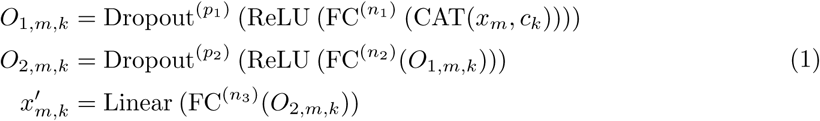

where *x′*_*m,k*_ represents the latent representation of the drug *m* with cell line *k*, which is the output of the *DSN*_*m*_. FC^(∗)^ represents a fully connected layer with * neurons and Dropout^(*p*)^ represents a dropout layer with probability *p*. Linear and ReLU stand for linear and rectified linear unit activation functions, respectively. CAT concatenates given features vectors.

#### Synergy Prediction Subnetwork (SPN)

SPN takes the concatenated vector embeddings calculated by *DSN*_*i*_ and *DSN*_*j*_ subnetworks. SPN is a three-layered FC network. Similar to DSN, we use RELU activation in the first two FC layers and linear activation at the output layer. Dropout, with a drop probability of 0.5, is applied after the first ReLU activation layer. SPN produces the predicted Loewe score of the drug pair < *i*, *j* > on cell line *k* denoted as *ŷ*_*i,j,k*_. The formulation of SPN can be summarized as follows:

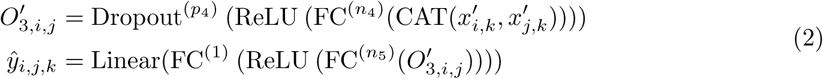

We train MatchMaker in an end-to-end fashion using a weighted version of MSE as the loss function (Equation 3). We modify the loss such that the examples that are more synergistic are weighted more. Our goal is to focus more on the examples that are highly synergistic because these are drug pair, cell line combinations with higher therapeutic potential. To achieve this, the weight of a trio in the loss calculation is set proportionally to its true Loewe score’s distance from the minimum of all Loewe scores in the training data (Equation 4).

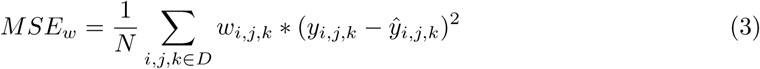

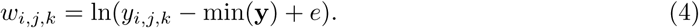

where *N* is the size of the training dataset *D* and *w*_*i,j,k*_ is the loss the weight for a trio (drug *i*, drug *j*, cell line *k*). **y** ∈ ℝ^*N*^ denotes the vector of true labels in the training set, *y*_*i,j,k*_ is the true Loewe score for the corresponding trio, *ŷ*_*i,j,k*_ is the predicted score and *e* is the Euler’s number.

### 2.5 Compared Methods

We compare the performance of MatchMaker with two available methods from the literature, which also exclusively use drug structure and the untreated gene expression profiles of the cell lines. DeepSynergy is a fully-connected multi-layered deep neural network based algorithm [16]. The second algorithm is TreeCombo, in which Janizek et al. use XGBoost algorithm [18] to predict the Loewe score of drug pair < *i*, *j* > in cell line *k*. Both methods input a concatenated vector of the drug chemical structure, and cell line features: *CAT* (*x*_*i*_, *x*_*j*_, *c*_*k*_). Note that this is different from MatchMaker architecture, which first learns an embedding per drug on the cell line of interest and then concatenates the embeddings to learn synergy score prediction.

We also compare MatchMaker with two other standard machine learning methods: (i) Random Forest, an ensemble method based on decision trees, and (ii) ElasticNet, a linear regression method that combines *L*_1_ and *L*_2_ regularization. To be consistent with the previous two methods, Random Forest [32] and ElasticNet [33] are also trained with the same concatenated input *CAT* (*x*_*i*_, *x*_*j*_, *c*_*k*_). Finally, we calculate a baseline model that for a drug pair-cell line trio, < *i, j, k* >, predicts the label as the mean of the synergy scores for all training examples that involve the cell line 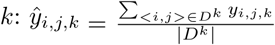 where *D*^*k*^ contains all experiments performed on cell line *k*.

## 3 Results

### 3.1 Experimental Setup

For performance assessment, we split the dataset into training, validation, and test folds with ratios 60%, 20%, and 20%, respectively. To ensure a fair comparison, we stratify the samples into folds using *leave-drug-combination-out approach* as described in Preuer et al.. In this approach, a drug pair appear in only either one of the train, validation, or test sets. We tune the parameters on the validation set and final performances are reported on the test set. All algorithms use the same sets for training, validation, and testing.

In addition to the above-mentioned setting, to assess the robustness of the results, we reevaluate MatchMaker using repeated 5-fold cross-validation. That is, we train the model with three folds, use the one fold for validation, and one fold for testing. We rotate these folds and additionally repeat this process 10 times with different random splitting of three folds. At the end of this, we obtain 50 MatchMaker models and 10 predictions per drug pair and cell line combination.

In tuning the DeepSynergy model parameters, we consider the best 10 parameter settings reported in Preuer et al. (see Table S2) and choose the best among them using the validation data. For the TreeCombo model, we use the parameter combinations provided in Janizek et al. (see Table S3). For Random Forest, we choose the best performing *maximum tree depth* ∈ {4, 6, 8, 10, 12} and *the number of trees* ∈ {10, 100, 500, 1000} parameters (Table S4). Finally, for ElasticNet *L1 ratio* ∈ {0.20, 0.40, 0.60, 0.80} and *α* ∈ {0.001, 0.01, 0.1, 1.0, 10, 100} parameters are considered (Table S5).

We train MatchMaker using early stopping mechanism [34]. That is, while minimizing the training loss, we monitor the validation loss and terminate the training if validation loss does not improve for *p* epochs (*p* = 100) and save the model that provided the best validation loss. We set the maximum number of epochs to 1000. We use the Adam optimizer [35] with the learning rate of 10−^4^ and batch size 128 to the loss function. We use Xavier initialization [36]. During training, a sample for a drug pair < *i*, *j* > is input to the network also in the reversed order, < *i*, *j* >. This is to train each DSN with every drug and to avoid order bias. During testing, we obtain predictions for both < *i*, *j* >, and < *i*, *j* > for all methods, and the final Loewe score is obtained by averaging these two predicted scores.

We train all the models on a SuperMicro SuperServer 4029GB-TRT with 2 Intel Xeon Golf 6140 Processors (2.3GHz, 24.75M cache) and 251GB RAM. MatchMaker and DeepSynergy are trained on 1 NVIDIA TITAN RTX GPU (24GB, 384Bit). The training times were approximately 5 hours for MatchMaker and 4 hours for DeepSynergy. We use Keras [37] with TensorFlow [38] backend for MatchMaker and DeepSynergy. We use scikit-learn [39] implementations for Random Forest and ElasticNet and the Python package of XGBoost to implement TreeCombo [18]. We train TreeCombo, ElasticNet and Random Forest on CPU, and their training times were approximately 1 hour, 30 mins and 5 hours, respectively.

### 3.2 Performance Comparison

#### Regression Performance

We compare the models’ predictive performances on the held out drug combinations using three performance metrics: (i) MSE, (ii) Pearson, and (iii) Spearman rank correlation coefficients between the actual Loewe scores and the predicted Loewe scores. Figure 3 summarizes the model performances for different algorithms and for different performance metrics. MatchMaker achieves the lowest MSE value, 79.49, while DeepSynergy is the next best performer with an MSE value of 112.6. TreeCombo’s predictions lead to 132.7 MSE. This corresponds to a 42% improvement over DeepSynergy and a 67% improvement over TreeCombo in terms of MSE. Of all the methods, Random Forest and ElasticNet perform worst and yield to the highest MSE values: 173.2 and 183.4, respectively. Note that the baseline model leads to 202.5 MSE. When models are evaluated in terms of the Pearson and Spearman rank correlation coefficients, a similar trend is observed. MatchMaker achieves the best performance in terms of Pearson, 0.79, and 0.74 Spearman correlation coefficients. Overall, the rankings of the models are the same across the three performance metrics and MatchMaker is consistently the best performing one among the five algorithms compared.

**Fig. 3:**
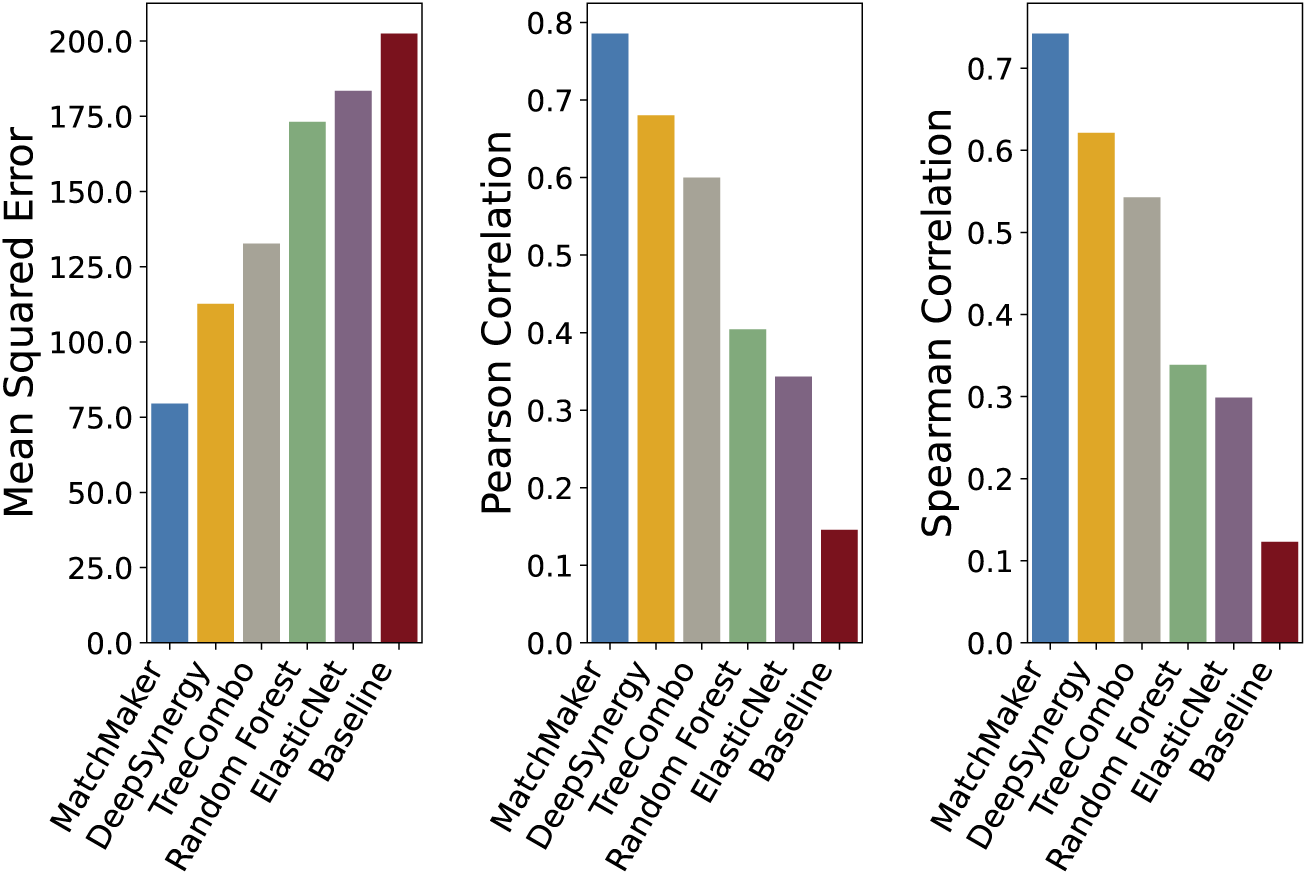
Performance comparison of models based on MSE, Pearson and Spearman correlation coefficients. MatchMaker achieved the best performances with 79.49 MSE, 0.79 Pearson and 0.74 Spearman’s correlation coefficients.

Next, we investigate if cell line gene expression data bring predictive power. MatchMaker trained in the same experimental settings but without the cell line gene expression features leads to an MSE of 131.9, Pearson correlation of 0.60, and Spearman correlation of 0.53. This clearly shows that cell line gene expression data is an important ingredient for the model, resulting in a substantial performance gain.

#### Classification Performance

We further assess the trained regression model’s ability to distinguish synergistic drug combinations and mimic a classification task for evaluation. That is, we binarize the actual synergy scores (*y* ∈ ℝ) such that those larger than a positive Loewe score threshold, *t*, constitute the positive class (synergistic) pairs 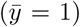. In contrast, those less than −*t* constitute the negative class 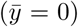, where *t* ∈ (0, 10, 20). We evaluate the predictive performance of the models on this binary classification task by the area under the receiver operating characteristics (ROC) curve. Figure 4 shows ROC curves for the different models when *t* is set to 10. Based on AUC, partial AUC (false positive rate up to 0.1; pAUC) and area under the precision recall curve (AUPR), MatchMaker outperforms the other compared methods by a large margin (see Table 1).

**Table 1:**
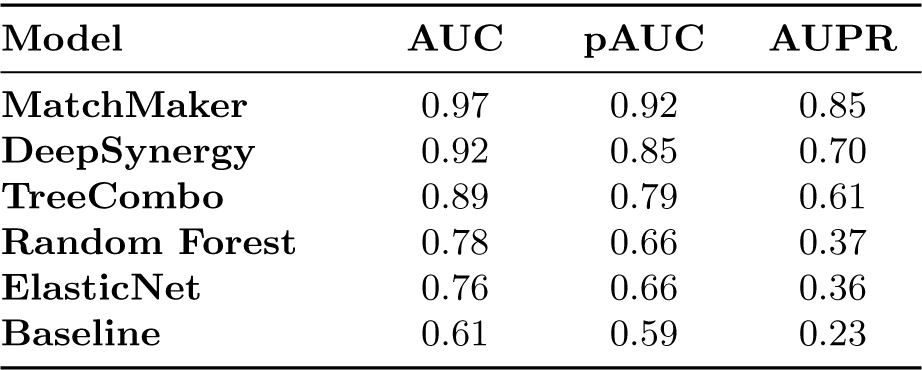
Performance comparison of the methods in the classification task. Area under ROC curve (AUC), partial AUC calculated up to false positive rate 0.1 (pAUC), and area under precision-recall curve (AUPR) are shown for all methods considered. Experiments with a Loewe score higher than 10 were classified as positive (synergistic) samples and those below −10 are classified as negative (antagonistic) samples.

**Fig. 4:**
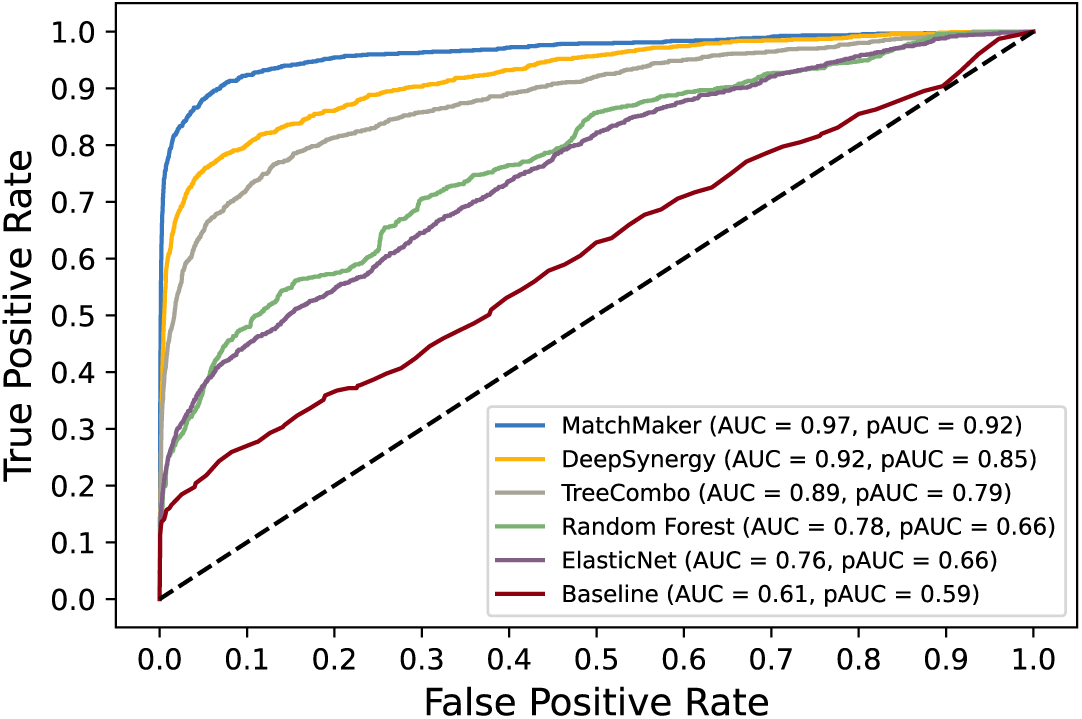
Receiver Operating Characteristics (ROC) curves for all methods in classification task. Synergistic combinations with a Loewe score higher than 10 were classified as positive samples and those below −10 are classified as negative. Corresponding AUC values and partial AUC (pAUC) values up to 0.1 FPR are also noted.

The same classification success holds when *t* is varied (see Figure S2 and S3; and Table S6 and S7). When *t* is set to 20 instead of 10, performances for all models increases as distinguishing very synergistic and very antagonistic examples is a relatively easier classification task. When *t* is set to 0, we are testing whether the model is able to differentiate the positive and negative Loewe scores (Figure S2 and Table S6). This is a more difficult task. Also note that, in this case, the definitions of the true positive and true negative are clouded by noise, as a Loewe score just barely above zero is not a very strong indicator of synergy. Even though the performances for all the models decrease when *t* = 0, MatchMaker still outperforms other methods in terms of the classification performance metrics.

The ultimate goal of in-silico synergy prediction models is to prioritize the candidates to be validated experimentally. Therefore, it is relatively more important to achieve higher performance at the top candidates with the highest predicted synergy scores. We also compare the models in terms of the pAUC score which considers AUC in a restricted false positive rate (FPR) range. When we compare the methods using pAUC up to 10% FPR, we observe that MatchMaker pAUC improvement is even larger than the pAUC improvement of the second best method. MatchMaker achieves 0.92 pAUC score, whereas DeepSynergy achieves 0.85 when *t* is set to 10. The same trend holds when *t* is set to 0 and 20 as well (Figure S2 and S3; and Table S6 and S7).

### 3.3 MatchMaker Performance Across Cell Lines and Drug Pairs

To understand the extent of variability of the model performance across cell lines, we first calculate the cell line specific performance metrics. That is, we measure the performance for all experiments performed on a cell line *k*. Figure 5 shows that there are few cell lines where the model performs poorly. The cell lines with very high MSE values are T98G, L-1236 and NCIH2122. We inspect the reason that could explain this performance difference. Yet, we do not see a clear association to the number of training examples available (1036, 1391, and 290, respectively) or their tissue types (brain, haematopoietic/lymphoid, and lung, respectively). Interestingly, when we analyze the consistently-predicted false positives (i.e., novel predictions), many experiments on T98G cell line are predicted to be false positive with a large MSE gap (see Section 3.4). Thus, one possibility is that the actual labels for these cell lines could be noisy and there is a systematic error on these cell lines’ experiments. There are also a few cell lines for which the model performs remarkably well. For the top 6 performing cell lines, the MSE is below 30. The best performing three cell lines are RD, A2780 and A427. When we inspect the correlation metrics, we observe that for 75 of the 81 cell lines, the Pearson correlation coefficient is above 0.7, indicating that for most of the cell lines MatchMaker performance is good. Finally, we observe for 71 of the 81 cell lines, the Spearman rank correlation coefficient is above 0.7. We provide these cell line specific evaluations in Figure S1 and Table S8.

**Fig. 5:**
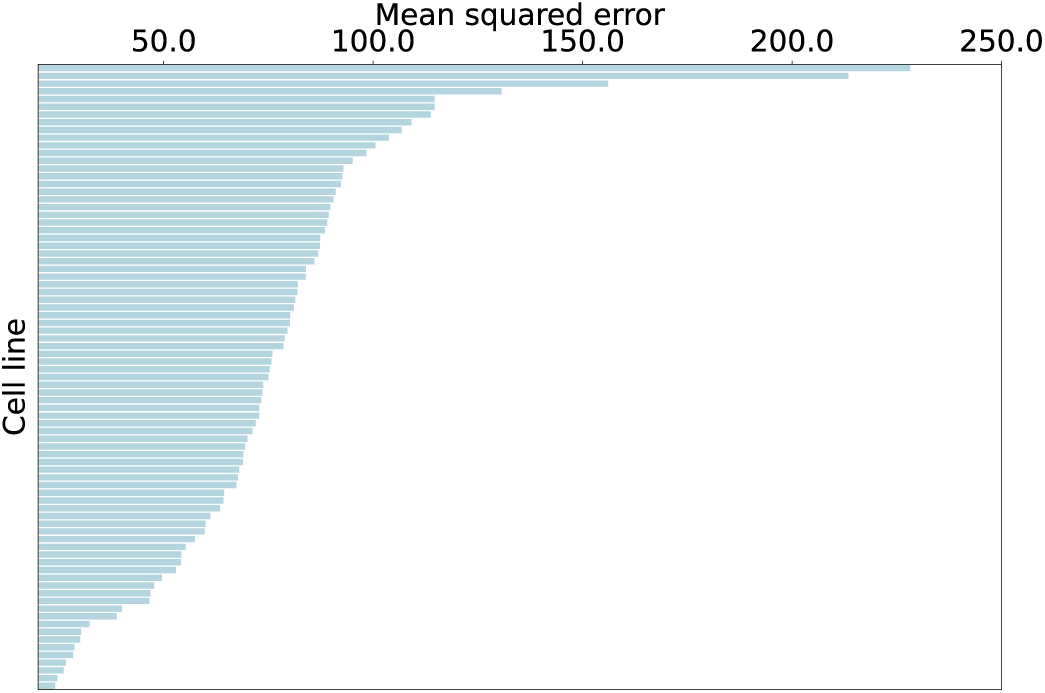
Cell line specific MSE performance of MatchMaker is shown.

Next, we evaluate the drug-pair specific predictive performance of MatchMaker. We consider the drug pairs that appear with more than one cell line in the drug pair-cell line combinations in the test data. Table 2 displays the top 10 drug pairs that yield the minimum MSEs. Over the 1082 unique drug pairs, we observe that 74% of them are below the overall MSE of 79.49. In terms of the correlation, we observe 46% and 35% of drug pairs are above 0.7 for Pearson and Spearman correlation coefficients, respectively. Thus, in terms of correlation metrics, the performance with respect to drug pairs vary more compared to performance with with respect to the cell lines. We see that all but one top-10 drug pairs have over 50 experiments in the test dataset. The average number of experiments (i.e., number of cell lines investigated) per drug pair is 25. This shows that the model performance overall benefits from large number of training samples.

**Table 2:**
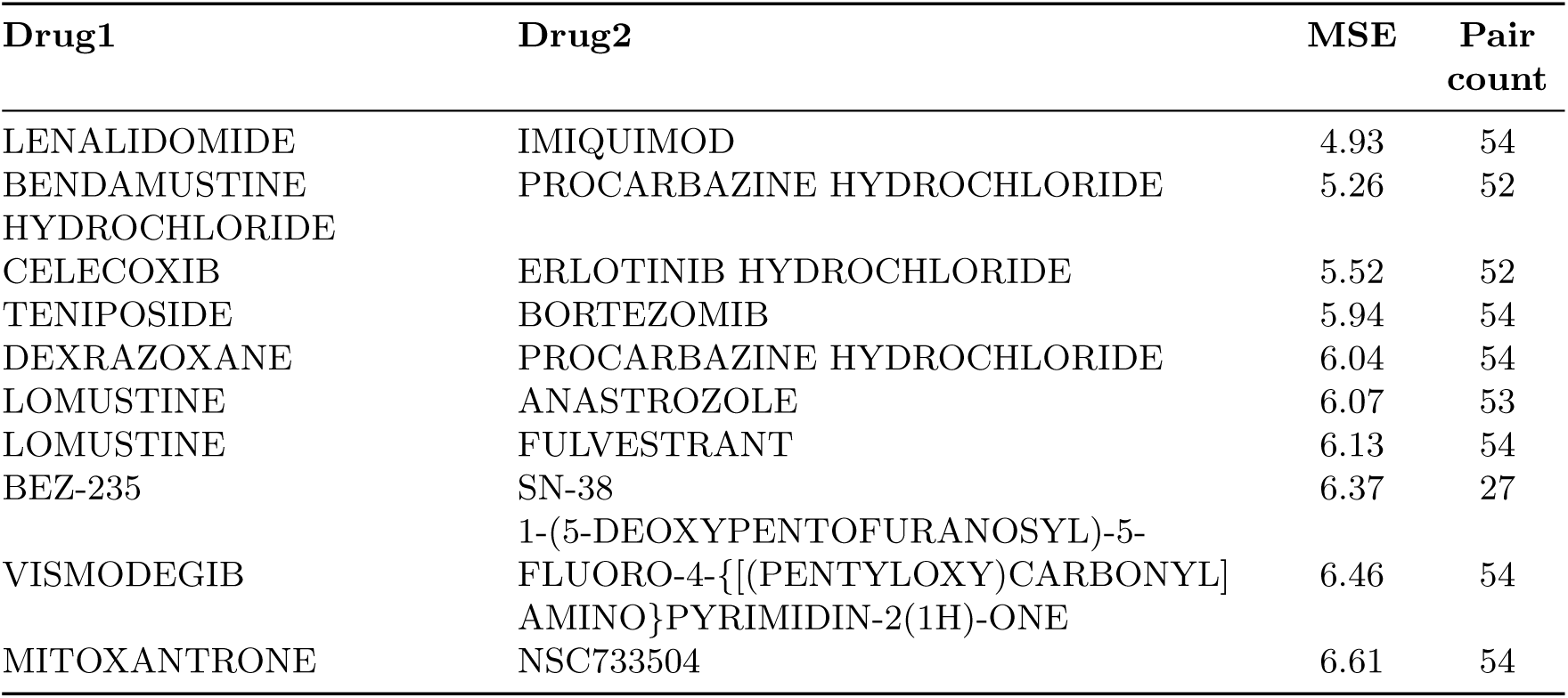
The top predictive drug pairs based on average MSE.

### 3.4 Novel Predictions

The ground truth labels for experiments in DrugComb could be noisy or could have systematic bias. Here, we check the drug-cell line combinations that are consistently predicted as false positives by MatchMaker. That is, the model predicts a particular drug pair in a cell line to interact in a synergistic fashion, where indeed it is labeled as antagonistic in the ground truth.

We find the trios that are consistently predicted as false positive in all 10 random initialized MatchMaker models. Note that they are in the test set 10 times despite we employ a 5-fold cross validation scheme as described in Section 3.1. Out of the 286, 421 combinations, we find 2, 540 drug pair-cell line combinations that are consistently marked as false positives. To get the ones that are more confident, we retain the combinations with actual Loewe score less than −10 and predicted Loewe score larger than 10. This leaves us with 25 combinations, which are provided in Table 3. We present the whole list of 2, 540 drug pairs in Table S9.

**Table 3:**
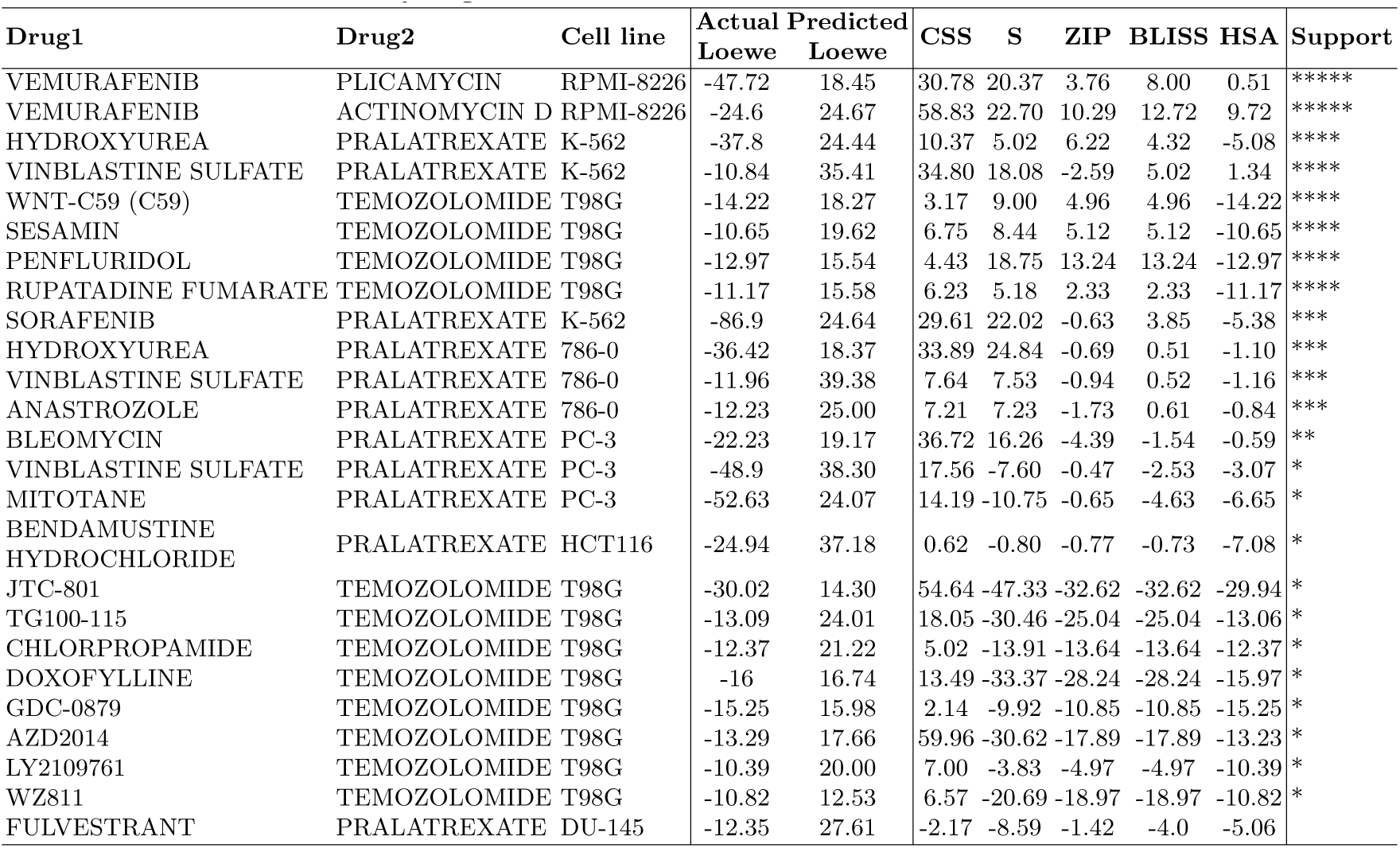
The list of confident false positives which are designated as novel synergistic drug pair predictions. The pairs that have actual Loewe score less than −10 and have MatchMaker -predicted Loewe score larger than +10 in all 10 random initializations of the model are listed. The subsequent five columns of the table list alternative drug sensitivity (CSS) and drug combinations sensitivity scores (S, ZIP, BLISS, HSA). The last four sensitivity scores are obtained from DrugComb database. The last column summarizes the support of these alternative measures. That is, the number of stars indicates the number of alternative measures that also indicate a synergistic interaction.

For these 25 trios, we also inspect drug combination sensitivity scores (CSS), and the other synergy measures: Bliss independence (BLISS) [40], Highest single agent (HSA) [41] and Zero interaction potency (ZIP) [42] and S synergy score [43]. For the two of the 25 trios, all of these measures indicate a synergistic interaction, while for the rest of the trios, there is a varying degree of support. We list this extra information in Table 3.

The first two trios in Table 3 are: <VEMURAFENIB, PLICAMYCIN> and <VEMURAFENIB, ACTINOMYCIN D> tested on RPMI-8226 cell line (haematopoietic/lymphoid tissue). For these pairs, (i) MatchMaker predicts a synergistic relationship, (ii) Loewe score indicates an antigonistic interaction, and finally, (iii) all other measures are in favor of synergysm. For one of these pairs we indeed find evidence in the literature: VEMURAFENIB and ACTINOMYCIN D interacts and VEMURAFENIB can increase the serum concentration of ACTINOMYCIN D [44].

The following six combinations in Table 3 are supported by at least three out of four synergy measures and by a positive CSS score. Among them, the PENFLURIDOL and TEMOZOLOMIDE (TMZ) combination has been shown to prolong the survival by (i) decreasing the tumor growth [45], and (ii) increasing the apoptosis in Glioblastoma (GBM) tumors [46]. The drug pair reduces the MGMT and GLI1 expression in T98G cell line along with several other GBM cell lines [46]. MatchMaker is also in disagreement with the actual truth Loewe scores in combinations of TMZ with (i) TG100-115, (ii) CHLORPROPAMIDE, and (iii) LY2109761. We find evidence in the literature supporting the effectiveness of these drug combinations. Li et al. state that TMZ enhances the effect of TG100-115 on GBM cells and improves survival in mice [48]. Zhang et al. show that the combination of LY2109761 and TMZ suppresses the tumor growth on T98G cell line. Finally, we find that CHLORPROPAMIDE can increase the serum concentration of TMZ [44].

Table 3 shows that for two drugs the difference between the predicted and the actual synergy scores are off by a large margin for many cases: (i) PRALATREXATE and (ii) TMZ. Moreover, these experiments are frequently performed on the T98G cell line. T98G is one of the cell lines on which MatchMaker performs the worst on average (see Figure 5). As discussed in Section 3.3, systematic error on this set of T98G experiments could be an explanation for these contradictory predictions.

## 4 Discussion

In this study, for the first time, we made use of the DrugComb dataset [20], which contains close to half a million drug combination experiments. This is an invaluable source for training complex models. We observe that the large dataset size and a new architecture that learns drug specific weights enable MatchMaker to improve the prediction power of the state-of-the-art approaches substantially. We investigate the benefit of having more training examples for a drug. We observe that drugs that have more than 100 training samples in the training data have an average MSE of 79.43, Pearson correlation coefficient of 0.79, and Spearman correlation coefficient of 0.79. This is in contrast to the drugs with less than 100 training samples which have MSE of 161.69, Pearson correlation coefficient of 0.64, and Spearman correlation coefficient of 0.68. We foresee that with the growing experimental dataset sizes will enable more complex architectures to be trained and pave the way to more precise drug synergy predictions. Although we have a very large dataset for training today, the number of possible combinations to exhaustively validate experimentally is still intractable. This means accurate and efficient computational prioritization of novel synergistic drug pair candidates will keep being an important research area.

In MatchMaker, we use only the drug chemical structure as the primary feature, which has been used in parallel tasks such as drug target identification [50] or drug side effect prediction [51, 52]. We also use the cell line specific gene expression profile to capture the context of the experiment. These features are selected on the basis of their general availability to ensure the model’s broad applicability to different drug combinations. The model presented here can be enhanced by integrating other rich biological information available on drugs and their targets. It has been previously shown that network-based approaches also provide powerful predictions [53]. Pathways and protein-interaction networks could be leveraged. However, when drug targets are not one of the nodes of these networks, it reduces the size of the training data that can be used. Despite this limitation, using this information without sacrificing a large amount of training data is a future work to be explored. Other than drug side effects, drug categorizations and drug-disease relationships can be further utilized to improve the performance of MatchMaker.

Loewe score was our choice of target label due to its widespread use in the literature for this task. However, there are other synergy measures that differ in the reference model of non-interaction such as BLISS, HSA, ZIP and S synergy score. It is straightforward to modify our framework to work with these alternative measures of synergy. The model can also be enhanced to predict all measures simultaneously in a multi-task learning setting, which will be explored in future work.

During the preprocessing of the drug combination dataset, we filtered out drug pairs for which the replicate experiments exhibit large disagreement on the biological activity of the drug pair. However, there were no replicate experiments for a large portion of the dataset (81%). This could be a limitation of the dataset, which might have introduced label noise in the training and evaluation processes. Another limitation is that the DrugComb dataset considers only cancer cell lines. Although the framework is applicable to other conditions and diseases, the model trained is inevitably biased towards cancer. The model still could serve as a pretrained model and be fine-tuned for other cell lines and drugs for other diseases.

## 5 Conclusion

Combination therapy is of great interest in drug development due to improved efficacy and reduced side effects. In this work, we present MatchMaker, a deep learning-based model that utilizes drug chemical structure and cell line specific expression patterns to predict if two drugs work synergistically. We trained this model on the largest experimental dataset released to date. Our results demonstrate that MatchMaker outperforms state-of-the-art approaches in terms of predicting the Loewe score. By pruning the otherwise unmanageable search space, MatchMaker provides a useful tool for prescreening and prioritizing the candidate drug pairs in silico.

## Acknowledgments

The authors would like to acknowledge the efforts of Mert Albaba for the investigation of the datasets. This study was supported in part by TUBA – GEBIP 2017 award to AEC. OT and AEC acknowledge supports from Bilim Akademisi BAGEP awards.

