## Supplementary File for "MatchMaker: A Deep Learning Framework for Drug Synergy Prediction" (from another version of this preprint: version 1, 2020-05-27)

No Author Given

No Institute Given

#### 1 Chemical Features

Drug chemical structure features are calculated by a Python library called *ChemoPy*. The chemical features includes 11 different chemical descriptor types with a total of 541 different features. Table S1 shows the chemical feature types and corresponding number of features.

**Table S1:** Drug chemical structure features collected from ChemoPy

| Chemical Feature Type | Feature Count |
|---|---|
| Constitutional descriptors | 30 |
| Connectivity descriptors | 44 |
| Kappa descriptors | 7 |
| Basak descriptors | 21 |
| E-state descriptors | 245 |
| Burden descriptors | 32 |
| Moreau-Broto autocorrelation descriptors | 32 |
| Moran autocorrelation descriptors | 32 |
| Geary autocorrelation descriptors | 32 |
| Molecular property descriptors | 6 |
| MOE-type descriptors | 60 |

#### 2 Hyperparameter Tuning

The hyperparameters of all methods in paper were optimized on the validation pairs and performance is measured on test pairs. The hyperparameters with their corresponding ranges for baseline methods are given at Table S2, S3, S4 and S5. The hyperparameter setting in Table S2 are taken from *Preuer et al.*

**Table S2:** Hyperparameters for DeepSynergy. The input data is normalized either by *tanh_norm* or *tanh* as given at *Preuer et al.*. *tanh* first standardizes and then applies hyperbolic tangent. *tanh_norm* first standardizes, applies hyperbolic tangent and standardizes again. The number of neurons in layers of DeepSynergy are separated with underscore character '_'

| Layers | Dropout Input | Dropout | Learning Rate | Preprocessing |
|---|---|---|---|---|
| 8182_4096_1 | 0.5 | 0.2 | 0.00001 | tanh_norm |
| 8182_8182_1 | 0.5 | 0.2 | 0.00001 | tanh_norm |
| 8182_2048_1 | 0.5 | 0.2 | 0.00001 | tanh_norm |
| 4096_4096_1 | 0.5 | 0.2 | 0.0001 | tanh |
| 2048_2048_1 | 0.5 | 0.2 | 0.0001 | tanh |
| 4096_2048_1 | 0.5 | 0.2 | 0.0001 | tanh |
| 4096_2048_1 | 0.5 | 0.2 | 0.0001 | tanh_norm |
| 8182_8182_8182_1 | 0.5 | 0.2 | 0.00001 | tanh_norm |
| 8182_4096_2048_1 | 0.5 | 0.2 | 0.00001 | tanh |
| 4096_4096_1 | 0.5 | 0.2 | 0.0001 | tanh_norm |

**Table S3:** Hyperparameters for XGBoost. The input data is normalized with *tanh_norm*. The number of estimator parameter is set to 1000 as given at *Janizek et al.* Maximum tree depth and learning rate ranges used same as *Janizek et al.*

| **Hyperparameter** | **Tuned values** |
|---|---|
| preprocessing | tanh_norm |
| maximum tree depth | 4, 6, 8, 10, 12 |
| learning rate | 0.05, 0.10, 0.15 |
| The number of estimators | 1000 |

**Table S4:** Hyperparameters for Random Forest. The input data is normalized with *tanh_norm*.

| **Hyperparameter** | **Tuned values** |
|---|---|
| preprocessing | tanh_norm |
| maximum tree dept | 4, 6, 8, 10, 12 |
| the number of estimator | 10, 100, 500, 1000 |

**Table S5:** Hyperparameters for ElasticNet. The input data is normalized with *tanh_norm*.

| **Hyperparameter** | **Tuned values** |
|---|---|
| preprocessing | tanh_norm |
| a | 0.001, 0.01, 0.1, 1.0, 10, 100 |
| L1 ratio | 0.20, 0.40, 0.60, 0.80 |

### 3 Prediction Performance

#### 3.1 Predictive performance on cell lines

We analyze the performance of MatchMaker on each cell line and show them at Figure S1

**Fig. S1:** Cell line specific performances. The figures show the (a) MSE, (b) Pearson and (c) Spearman correlation performances for each cell line.

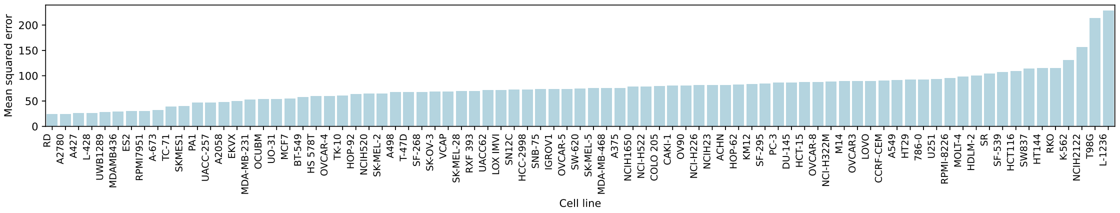


**(a)**

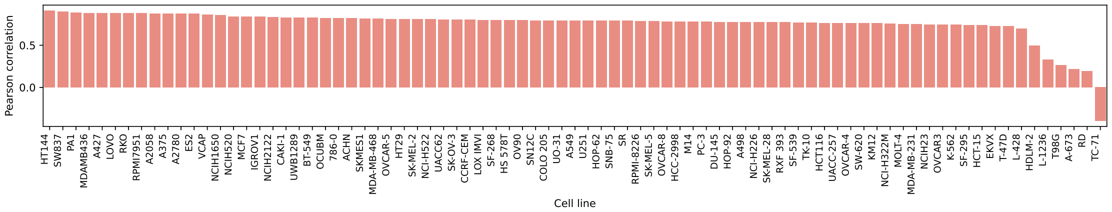


**(b)**

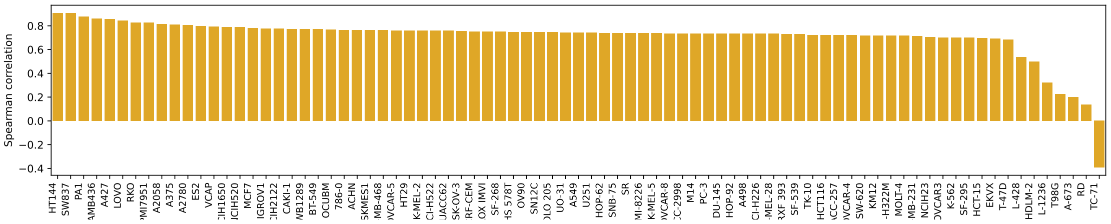


**(c)**

##### 3.2 Predictive performance on novel drug combinations

In addition to the classification results shown the main manuscript with the threshold $t = 10$, we further used $t$ as 0 and 20 to binarize the synergy score labels to classes and provide Area Under Receiver operating characteristics (ROC) curves at Figures S2 and S3.

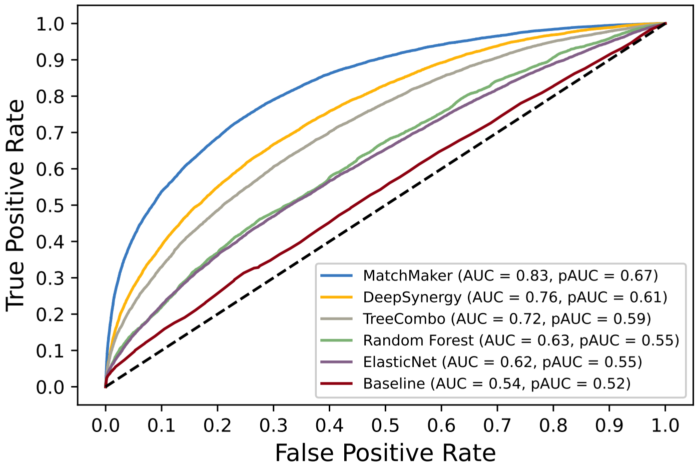


**Fig. S2:** Area Under Receiver operating characteristics (ROC) curves for all methods in classification task where threshold is 0.

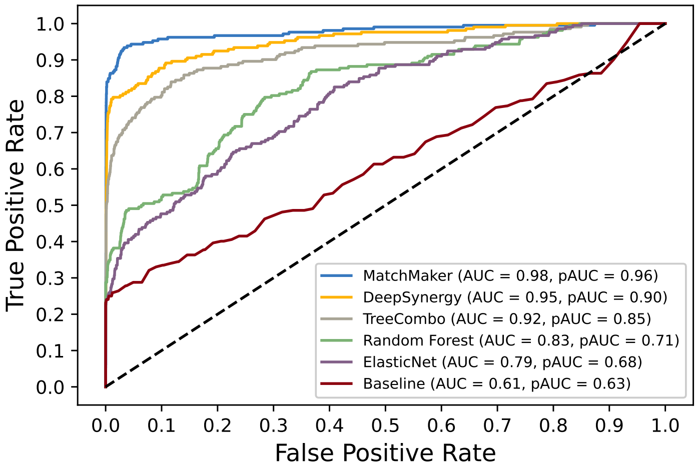


**Fig. S3:** Area Under Receiver operating characteristics (ROC) curves for all methods in classification task where threshold is 20.

**Table S6:** Performance comparison of the methods in the classification task. Area under ROC curve (AUC), partial AUC calculated up to false positive rate 0.1 (pAUC), and area under precision-recall curve (AUPR) are shown for all methods considered. Experiments with a Loewe score higher than 10 were classified as positive (synergistic) samples and those below -10 are classified as negative (antagonistic) samples.

| Model | AUC | pAUC | AUC PR |
|---|---|---|---|
| **MatchMaker** | 0.83 | 0.67 | 0.65 |
| **DeepSynergy** | 0.76 | 0.61 | 0.52 |
| **TreeCombo** | 0.72 | 0.59 | 0.48 |
| **Random Forest** | 0.63 | 0.55 | 0.38 |
| **ElasticNet** | 0.62 | 0.55 | 0.37 |
| **Baseline** | 0.54 | 0.52 | 0.30 |

**Table S7:** Performance comparison of the methods in the classification task. Area under ROC curve (AUC ROC), partial AUC calculated up to false positive rate 0.1 (pAUC ROC), area under precision-recall curve (AUC PR) are shown for all methods considered. Experiments with a Loewe score higher than 20 were classified as positive (synergistic) samples and those below -20 are classified as negative (antagonistic) samples.

| Model | AUC | pAUC | AUC PR |
|---|---|---|---|
| **MatchMaker** | 0.98 | 0.96 | 0.85 |
| **DeepSynergy** | 0.95 | 0.90 | 0.75 |
| **TreeCombo** | 0.92 | 0.85 | 0.65 |
| **Random Forest** | 0.83 | 0.71 | 0.41 |
| **ElasticNet** | 0.79 | 0.68 | 0.33 |
| **Baseline** | 0.61 | 0.63 | 0.28 |

##### 3.3 Confidence on predictive performance of MatchMaker

In addition to main results, we also assess the confidence of the performance of MatchMaker . The data is splitted into 5 fold by leave drug combination out approach. Each time, MatchMaker is trained on 3 training fold and validated on 1 validation fold. Then, performance is tested with the remaining 1 test fold. For each test fold, MatchMaker is trained 10 times with random initialization which attains 50 runs in total. Area under ROC curves are averaged over the 50 runs. In Figure S4, the average ROC curve is shown with solid line and shaded area represents the error bars with one standard deviation.

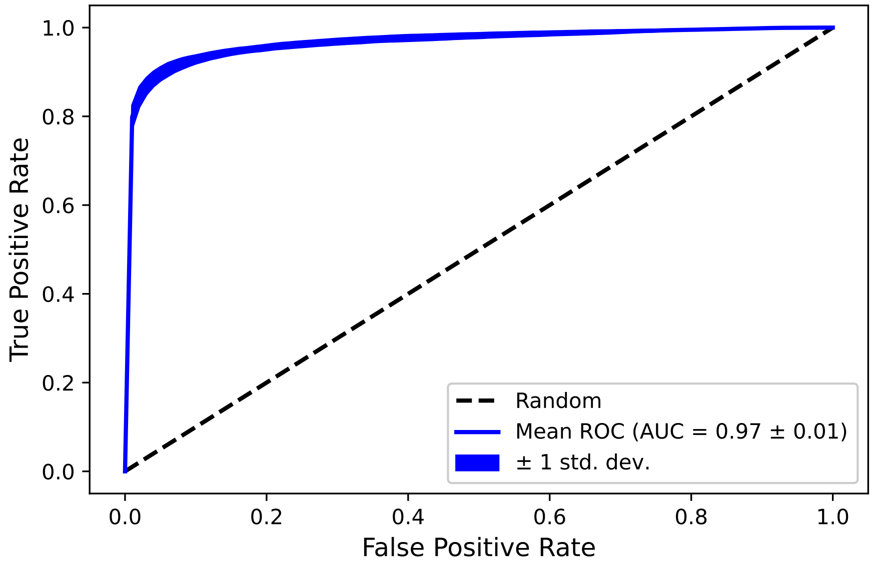


**Fig. S4:** Area Under Receiver operating characteristics (ROC) curve for MatchMaker and confidence interval in classification task where threshold is 10.
